# Quantifying photosynthetic restrictions

**DOI:** 10.1101/2024.05.24.595758

**Authors:** Chandra Bellasio

## Abstract

Quantifying the effect of factors controlling CO_2_ assimilation is crucial for understanding plant functions and developing strategies to improve productivity. Methods exist in numerous variants and produce various indicators, such as limitations, contributions, and sensitivity, often causing confusion. Simplifications and common mistakes lead to overrating the importance of diffusion – whether across stomata or the mesophyll. This work develops a consistent set of definitions that integrates all previous methods, offering a generalised framework for quantifying restrictions. Ten worked examples are provided in a free downloadable spreadsheet, demonstrating the simplicity and applicability to a wide range of questions.

## Introduction

Photosynthetic gas exchange routines measure CO_2_ uptake (assimilation, *A*) and water vapour release (transpiration, *E*), under a range of external CO_2_ concentration and light intensities (Busch et al. 2024). Raw data is first processed by the gas analyser software, to derive stomatal conductance to CO_2_ (*g*_S_), together with the CO_2_ concentration in the substomatal cavity (*C*_i_) using mass-balance and diffusion theory (von Caemmerer and Farquhar 1981). The resulting ordered pairs (*C*_i_, *A*) can be described by mathematical models of photosynthesis. These are typically adjusted to match data through curve fitting procedures, which return parameters that capture physical or biochemical characteristics of leaves (Bellasio et al. 2016b; Bellasio et al. 2016a).

A variety of models exist that are typically classified as mechanistic or empirical. In mechanistic models, expressions describe biochemical processes known to occur within leaves; in empirical models, expressions describe the relationship between data patterns. The distinction is merely artificial, because mechanistic models still rely on data, while empirical models describe patterns that ultimately depend on the same underlying processes. Mechanistic models are typically validated by direct measurements of physical or biochemical properties, and are modular (they can be modified to include more processes, and the description of processes can be refined to virtually any desired level of detail), allowing to accommodate a virtually unlimited range of real or hypothetical scenarios. Empirical models are generally simple, and may become unreliable when used in environmental conditions differing from those under which they were calibrated, but are typically more accurate for describing empirical datasets and valid for all photosynthetic types.

Additional analyses can quantify how physical or biochemical properties, or their experimental manipulation, may restrict assimilation. Restriction analyses seek to gain insights into the state of the assimilation process, rather than estimating the quantities defining it. Various methods based on different assumptions and conventions abound in the literature, and the lack of an objective criterion for their selection often renders the choice of methods “*to some extent arbitrary*” (Jones 1985). The aim of this study is to establish a framework with general validity that is straightforward to implement and unambiguous.

The theoretical basis of restriction analysis were laid by Gaastra (1959). Early studies made the erroneous assumption that CO_2_ concentration at the mesophyll carboxylating sites (*C*_M_) is zero. In the 1970s, biochemical CO_2_ uptake was treated as a linear function akin to Ohm’s Law of electrical resistance (Jarvis 1971; Jones 1973; Jones and Slatyer 1972; Prioul and Chartier 1977). This “*linear resistance analysis is in most cases invalid”* (Farquhar and Sharkey 1982) because photosynthesis typically responds non-linearly with a saturating response to CO_2_. In addition, rather than conveying information about the process, resistance confounds with a physical characteristic, making it difficult to relate to the sought process information. Consequently, I deem these methods undesirable and I will not address them further.

The concept of restriction can be related to that of mathematical sensitivity, for review see Jones (1985). Absolute and relative sensitivity (elasticity) quantify the extent to which various input variables (such as *C*_i_) exert control over the process of assimilation. Control analyses have the benefit that, once model parameterization is defined, outputs are unequivocal (Jones 1973). The drawback is that control coefficients are relatively complex to derive, often difficult relate to actual physiological phenomena, and are applicable only to infinitesimal change.

Extending sensitivity analysis to finite changes between two well-identifiable conditions of interest, for instance resulting from experimental manipulation, approximates the response of assimilation to CO_2_ concentration at the mesophyll carboxylating sites (*A*/*C*_M_ curve) with a straight line. This practice, which has been widely used both for C_3_ plants (Grassi and Magnani 2005) and in its C_4_ variant (Cano et al. 2019) introduces error (Deans et al. 2019) and is fundamentally mistaken (Buckley and Diaz-Espejo 2015). The solution proposed by Prioul et al. (1984) was to approximate the *A*/*C*_M_ curve with many small steps; however, the need to derive assimilation graphically and to process data manually, meant that most studies calculated assimilation in only three points. This raised another issue that the order in which these points are evaluated influences the result of the analysis (path dependency).

Recent key advancements in restriction analysis have been switching from a graphical to a computerised method where drawn curves were replaced by fitted empirical models (Bellasio et al. 2016b). That approach was later utilized to compare C_4_ and C_3_ plants but remained limited, as it still used the classical three-point calculation, and was only applied to stomatal and non-stomatal limitations (Bellasio et al. 2018). Buckley and Diaz-Espejo (2015) developed over the intuition of Prioul et al. (1984) by increasing the number of steps to any large number of infinitesimal transitions. Their method had the main benefits of being unambiguous, avoiding linearization of assimilatory responses over finite intervals, and implicitly containing sensitivity, thereby reconciling contribution with control analysis. However, it only applied to contribution analysis in C_3_ plants, leaving the need for a method that is universally applicable.

This study develops a unified theory and terminology that can be used to analyse controls, limitations and contributions in all plants, and shows the implementation of common procedures in spreadsheets freely available for download, demonstrating the framework’s simplicity and applicability to a wide range of questions.

### Theory: constructing a general framework

A first common approach called ‘limitation analysis’ involves comparing a healthy plant with its potential attainable under idealized conditions. The analysis quantifies the gain in assimilation achieved by sequentially ‘lifting’ limitations until photosynthesis reaches its maximum ‘limit’. Hypothesize a healthy plant photosynthesizing under ordinary operational conditions (Figure 1a). Limitation analysis simulates a transition along a hypothetical dimension that lifts all restrictions (number 1 in Figure 1a).

**Figure 1.**
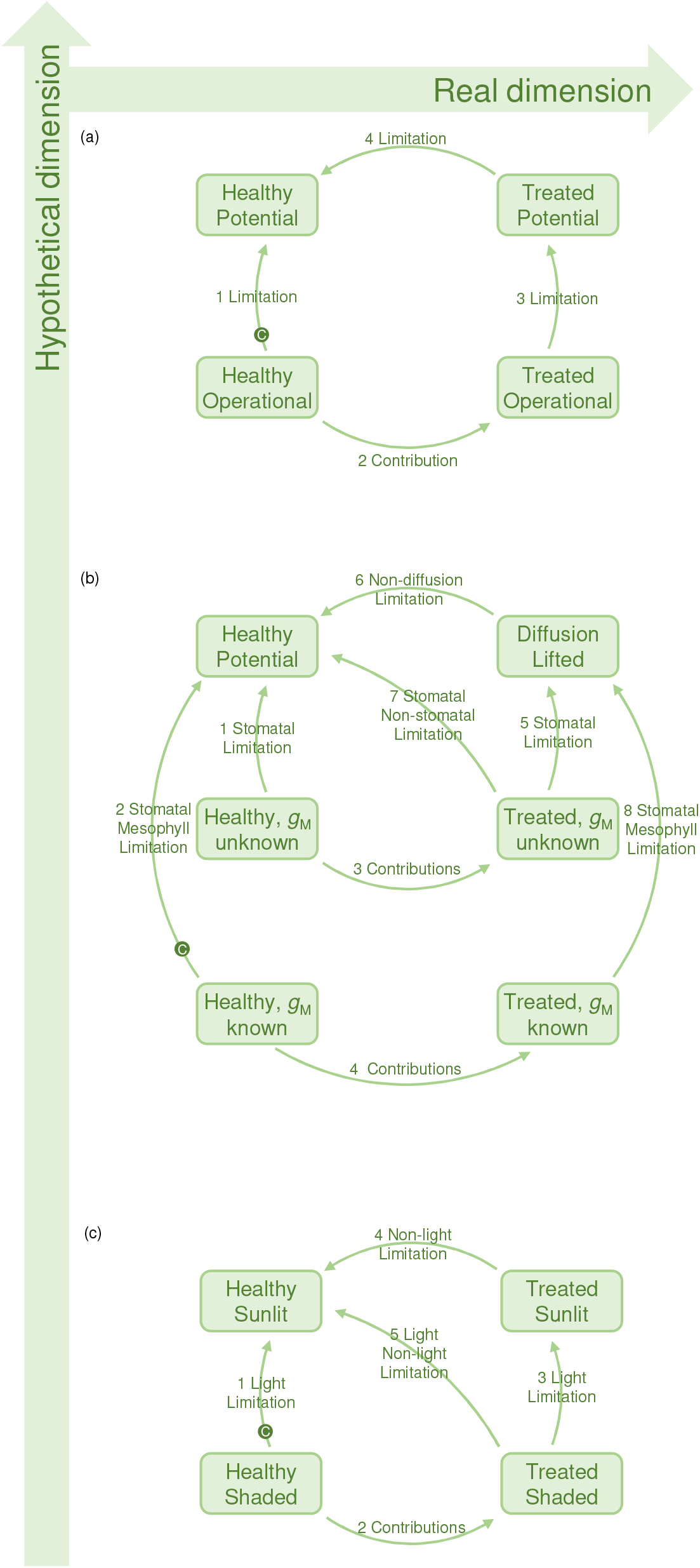
Generalised framework of restriction analysis. Restrictions may be evaluated along a real and an hypothetical dimensions. The analysis of limitations quantifies the restriction independently in each plant, compared their maximum potential. The analysis of contribution analyses the transition between two real conditions. The analysis of control (sensitivity and elasticity) can be done at any intermediate points of the transitions (exemplified by a ‘C’ in a green circle that may be swinging over all arrows). Panel (a) general framework; Panel (b) diffusional and non-diffusional restrictions; Panel (c) light and non-light restrictions.

An alternative approach called ‘contribution analysis’ compares a healthy plant with the same or a similar plant after the imposition of a treatment. The analysis attributes the changes in assimilation to the various quantities responsible for causing the change, that is, ‘contributing’ to that change in assimilation. Consider, for instance, a set of plants subjected to a generic treatment and left to photosynthesize under the same ordinary operational conditions as the healthy plants. Contribution analysis simulates the effect of the treatment, assigning the change in assimilation to the drivers causing it (number 2 in Figure 1a).

The strength of this new generalised framework lies in its ability to combine restriction analyses in any nuance. The extent to which treated plants are limited under ordinary conditions relative to their maximum limit, can be quantified through a limitation analysis (number 3 in Figure 1a). Any residual difference between the potential of the treated plant and that of the healthy plant can be evaluated through an idealised transition that reverts the treatment (number 4 in Figure 1a).

Analyses can also be combined. For instance, in a time-course experiment where a plant undergoes progressive treatment and is sampled at subsequent time points, limitation analysis can be performed at each time point, while contribution analysis can be used to compare these subsequent time points. Control analysis can be overlaid at any given intermediate point along transitions (a ‘C’ in Figure 1a) to determine the extent to which an infinitesimal change in the transition is controlled by environmental drivers and the underpinning photosynthetic characteristics.

To formalize this framework, it is necessary to assume that photosynthesis can be described mathematically. Let the function *f* be a set of expressions describing the dependence of any generic output variable Ω on a set of inputs, which are termed *a* (a generic input) and ***b*** (the set of all other inputs):

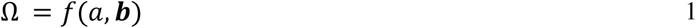

This is exemplified by the red line in Figure 2, which plots the values of Ω on the ordinate as a function of *a* on the abscissa, obtained for the specific input set ***b***_I_ (parameterization). Any situation with well-defined conditions, where *a* and ***b*** are known, will be referred to as a ‘state’. These may be the healthy and treatment operational conditions, or the hypothetical scenarios in which limitations are lifted. Only a few states are typically of interest, but, let the number of states be any integer *M*, with *m* being any generic state counted in Roman numbers (*m* = I, II, III … . *M*).

**Figure 2.**
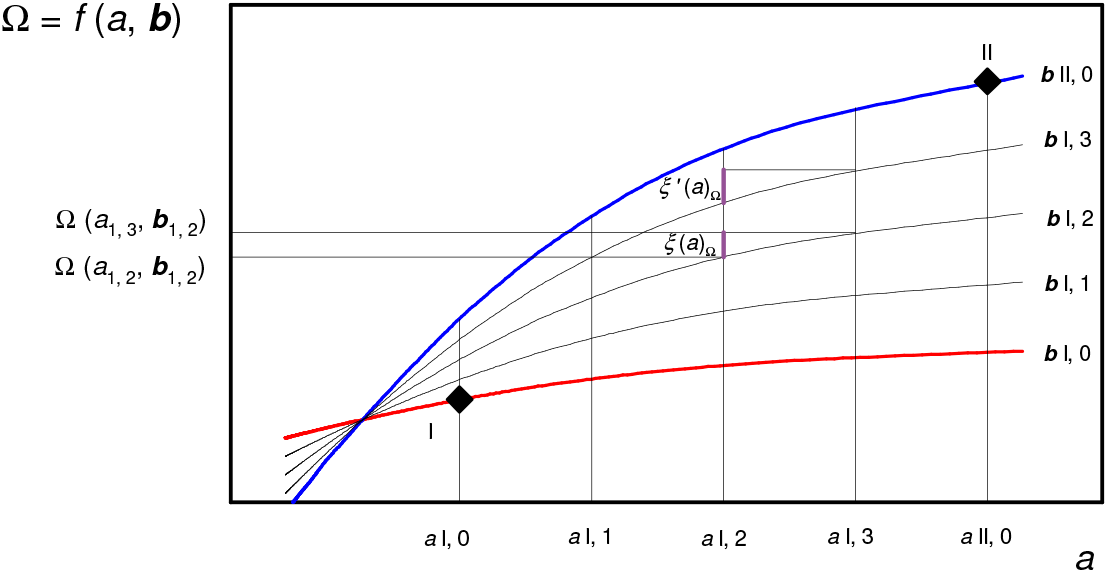
Schematic of the generalised framework of definitions. A generic output variable Ω is described through a mathematical model as a function of the variable *a* (in abscissa) and a set of other inputs ***b***. The conditions in which *a* and **b** are known are called ‘states’, here represented by two black diamonds. The function Ω = *f*(a, ***b***) parameterised with the known vector ***b***, for state I is shown in red, and for state II is shown in blue. The transition between the two known states is divided in four equal intervals, which are bordered by thin black lines. The value of the inputs *a* and ***b*** is assumed to vary linearly in the intervals. The marginal contribution *ξ* of the variable *a* to the change in the output Ω is shown in purple. The value of *ξ* ′ becomes equal to that of *ξ* for large values of *K*.

Any transition between two states is divided in a number of intervals *K*, being the total number of intervals, with *k* being any generic intermediate interval counted in integers (*k* = 0, 1, 2, 3 … . *K*). In Figure 2, the transition between state I and state II is divided in *K* = 4 intervals.

Following Buckley and Diaz-Espejo (2015) the value of a generic input in the interval *k* scales linearly between the values in the two adjacent known states *m* and *m*+I as:

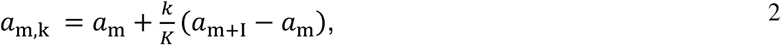

and the calculation is analogous for all other inputs in the set ***b***.

Within the interval *k* to *k*+1 the ‘marginal contribution of *a* to the change in Ω’ is defined as:

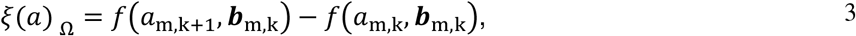

and is visualised in Figure 2 for the transition from *a*_I, 2_ to *a*_I, 3_. This definition of marginal contribution corresponds to that of Eqn 5b in Buckley and Diaz-Espejo (2015). Eqn 3 can be formulated in the analogous form *ξ*_’_(*a*)_Ω_ = *f*(*a*_m_,_k+1_, ***b***_m_,_k+1_) − *f*(*a*_m_,_k_, ***b***_m_,_k+1_) (shown in purple in Figure 2). Because *f*(*a*, b) is typically non-linear, *ξ*’(Ω)_a_ differs from *ξ*(Ω)_*a*_ by an error *ε*, see Deans et al. (2019). For instance, in Figure 2, *ξ*’(*a*_)Ω_ is 30% higher than *ξ*(*a*)_Ω_.*ε* decreases monotonically with *K*, therefore this issue vanishes when a suitably large value of *K* is chosen; in the examples provided *K*=1000 following Buckley and Diaz-Espejo (2015).

The key metric of contribution analysis is the ‘total contribution of *a* to the change in Ω’, which, in the transition from state I to state *M*, is the sum of the marginal contributions over all intermediate intervals and states:

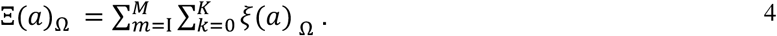

This definition of total contribution corresponds to that of Eqn 5c in Buckley and Diaz-Espejo (2015).

The ‘relative contribution of *a* to the change in Ω’ between the state I and the state M is:

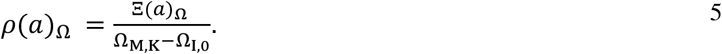

The definition of relative contribution corresponds to Eqn 25 in Jones (1985).

The key metric in limitation analysis is the ‘limitation by *a* to the change in Ω’, which, between the state I and the state M is:

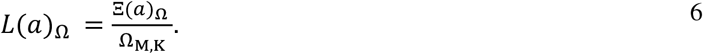

This definition of limitation corresponds to Eqn 13 in Farquhar and Sharkey (1982), Eqn 13 in Jones (1985), and to Eqn 7 in Buckley and Diaz-Espejo (2015) but expressed as percentage therein.

The key metrics in control analysis are absolute and relative sensitivity. The absolute ‘sensitivity of Ω to *a*’, calculated in a particular state and interval (*m, k*), is:

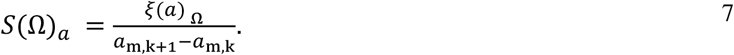

Sensitivity, a finite-difference estimate of 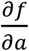, corresponds to Eqn 1 in Jones (1985) and to the partial derivatives in Eqn 1 of Buckley and Diaz-Espejo (2015).

Finally, the relative sensitivity better referred to as ‘elasticity of Ω to *a*’, calculated in a particular state (*m, k*), is:

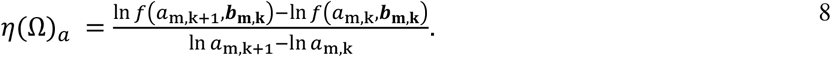

Elasticity, a finite-difference estimate of 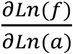 correspond to the relative limitation (*l*_S_, *l*_M_) in

Eqn 7 in Grassi and Magnani (2005), to the partial derivatives in Eqn 3 of Buckley and Diaz-Espejo (2015).

There will be as many analogous formulations of Eqns 1 to 8 as there are model inputs. All definitions given above are generally valid, but below I will only explore the case in which the output Ω is net photosynthetic CO_2_ uptake (*A*), and the function *f* is hence a model of photosynthesis.

### Applications

#### Diffusional and non-diffusional restrictions

Here I will initially use the empirical model of Prioul and Chartier (1977), because it is simple, suitable for all common restriction analyses, and can be used for C_3_ and C_4_ photosynthesis (Bellasio et al. 2018) as well as intermediate types alike. In the formulation of (Bellasio et al. 2016b) assimilation is:

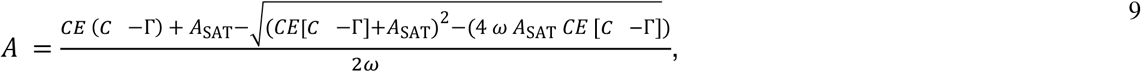

where, *A*_SAT_ is the CO_2_-saturated rate, *CE* is the initial slope, *C* may be *C*_i_ or *C*_M_, Γ is the *x*-intercept, *ω* is defining curvature.

CO_2_ enters the leaf through gaseous diffusion across stomata and the air passages between cells, dissolves in water, and diffuses in solution through the cell wall, plasmalemma and through the mesophyll to reach the Rubisco or phosphoenolpyruvate carboxylase sites of carboxylation, located in the chloroplasts of C_3_ leaves or in the cytosol of C_4_ leaves. The ease of this process is the total conductance 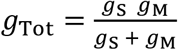, where mesophyll conductance (*g*_M_) compounds all processes that are not stomatal. The leaf supply function can be written as:

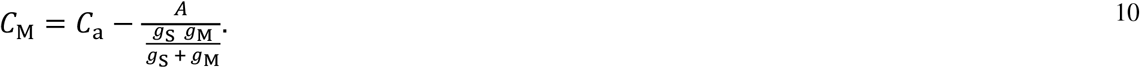

Combining Eqn 9 and Eqn 10 results in a quadratic for *A*:

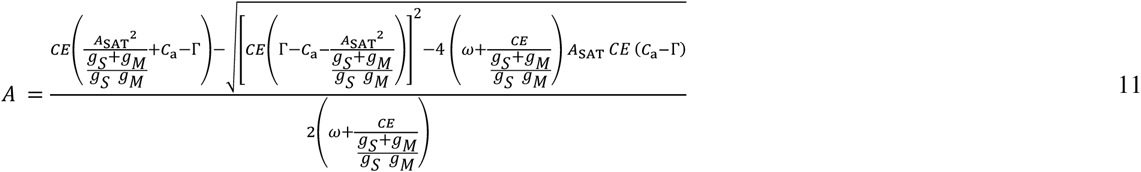

Eqn 11 will be used to evaluate diffusional and non-diffusional restrictions.

##### Stomatal limitation

Stomatal limitation is the reduction of assimilation due to stomatal closure that occurs in a leaf in a real, ordinary operational conditions as compared to a hypothetical case where CO_2_ would freely enter the leaf (the maximum limit of stomatal conductance is assumed to be infinity). The fixed characteristics of the plant *CE*, Γ, and *ω* (the vector ***b*** in Eqn 1) are found by curve fitting (procedures are available in Workbook I), that is, by inputting the measured value of *C*_i_ in Eqn 9 and adjusting their value until Eqn 9 outputs the output value of assimilation that is most similar to that measured. The values fitted to example data are shown in Table 2. Stomatal limitation is lifted in a single transition (number 1 in Figure 1b) by calculating Eqn 11 in all *K* intervals between state I where *g*_S_ equals *g*_Sop_, and state II, the hypothetical scenario where *g*_S_ is set to a high value (Procedures are available in Workbook II). A numerical example is shown in Figure 3A. The corresponding calculations are shown in Sheet 1 of Workbook III, where control coefficients (Eqn 7 and 8) are calculated for each step of the transition. The same procedure can be utilized to compare two known states resulting from any environmental perturbation (transition number 3 in Figure 1b), as long as they can be described by the same *A/C*_i_ response curve, simply by entering the two relevant *g*_S_ values.

**Table 1.** Acronyms, definitions, variables, and units used.

| Symbol | Definition | Units |
| --- | --- | --- |
| *a* | any generic input of a mathematical model |  |
| *A*, *A<sub>op</sub>* | Net assimilation, unspecified or measured under ordinary, operational conditions | $\mu\text{mol CO}_2 \text{ m}^{-2} \text{ s}^{-1}$ |
| *A<sub>SAT</sub>*, *A<sub>SAT</sub>'* | CO<sub>2</sub> saturated *A*, unspecified of after a generic treatment, respectively | $\mu\text{mol CO}_2 \text{ m}^{-2} \text{ s}^{-1}$ |
| ***b*** | a generic set of inputs of a mathematical model which includes all parameters and variables that are not *a* |  |
| *C<sub>a</sub>* | CO<sub>2</sub> concentration outside the leaf | $\mu\text{mol CO}_2 \text{ mol air}^{-1}$ |
| *CE*, *CE'* | Initial slope of the *A/C<sub>i</sub>* curve, unspecified of after a generic treatment, respectively | $\text{mol air m}^{-2} \text{ s}^{-1}$ |
| *C<sub>i</sub>*, *C<sub>iop</sub>* | CO<sub>2</sub> concentration in the substomatal cavity as calculated by the IRGA, unspecified, or under operational conditions | $\mu\text{mol CO}_2 \text{ mol air}^{-1}$ |
| *C<sub>M</sub>* | CO<sub>2</sub> concentration at the mesophyll carboxylating sites | $\mu\text{mol CO}_2 \text{ mol air}^{-1}$ |
| Contribution | Any change in an output variable which is due to a change only in one input, it can be marginal [small *csi*, $\xi$, Eqn 3], total [capital *csi*, $\Xi$, Eqn 4], or relative [*eta*, $\eta$, Eqn 5]. | $\mu\text{mol m}^{-2} \text{ s}^{-1}$, the relative is dimensionless |
| *E* | Leaf level water transpiration rate | $\text{mmol H}_2\text{O m}^{-2} \text{ s}^{-1}$ |
| *g<sub>BS</sub>* | Bundle sheath conductance to CO<sub>2</sub> diffusion | $\text{mol m}^{-2} \text{ s}^{-1}$ |
| *g<sub>M</sub>* | Mesophyll conductance to CO<sub>2</sub> diffusion | $\text{mol air m}^{-2} \text{ s}^{-1}$ |
| *g<sub>S</sub>* | Stomatal conductance, here to CO<sub>2</sub> diffusion also when unspecified, elsewhere often referred to as *g<sub>sc</sub>* | $\text{mol air m}^{-2} \text{ s}^{-1}$ |
| *g<sub>Tot</sub>* | Total conductance to CO<sub>2</sub> diffusion, the reciprocal of *g<sub>Tot</sub>* is the sum of the reciprocal of *g<sub>S</sub>* plus the reciprocal of *g<sub>M</sub>* | $\mu\text{mol m}^{-2} \text{ s}^{-1}$ |
| *k*, *K* | Any generic interval between states and the total number of intervals, respectively |  |
| Limitation | The total contribution divided by the maximum value of the function in the interval, Eqn 6 |  |
| *L<sub>NS</sub>* | Non-stomatal limitation to photosynthesis | dimensionless |
| *L<sub>S</sub>* | Stomatal limitation to photosynthesis | dimensionless |
| *m*, *M* | Any generic state, and the total number of states, respectively |  |
| Restriction | Any generic hindrance to photosynthetic activity |  |
| Sensitivity | The marginal contribution divided by the value of the variable, relative to the change in input divided by the value of the input, Eqn 7 |  |
| State | Any condition in which a photosynthetic model can be defined (by knowing *a* and ***b***) |  |
| $\mu$ | *A<sub>SAT</sub>* divided by *CE* | |
| $\Omega$ | Any generic output of a mathematical model, a function of *a* and ***b*** | |
| $\omega$ | Curvature of the non-rectangular hyperbola describing the *C<sub>i</sub>* dependence of *A* | dimensionless |

**Table 2.**
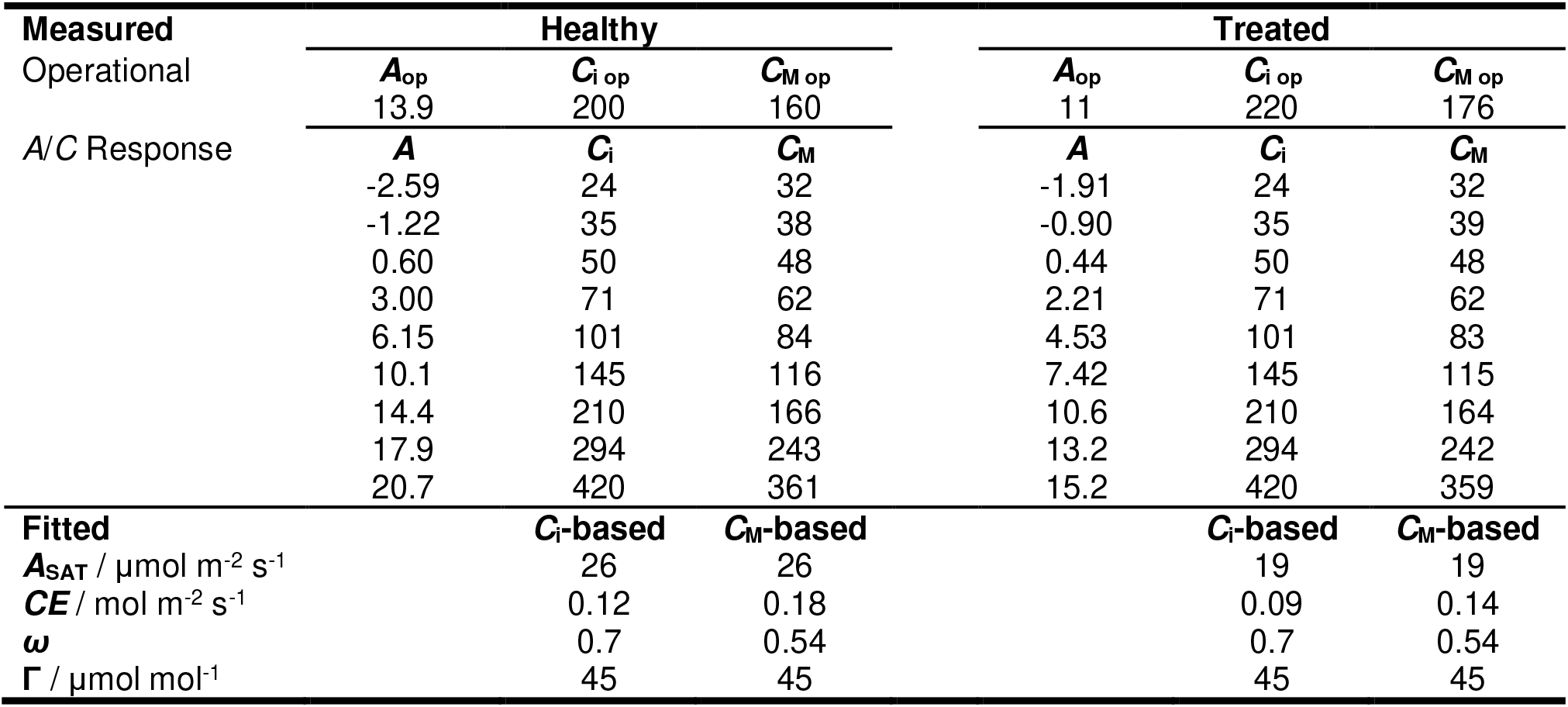
Synthetic data used in the examples.

**Figure 3.**
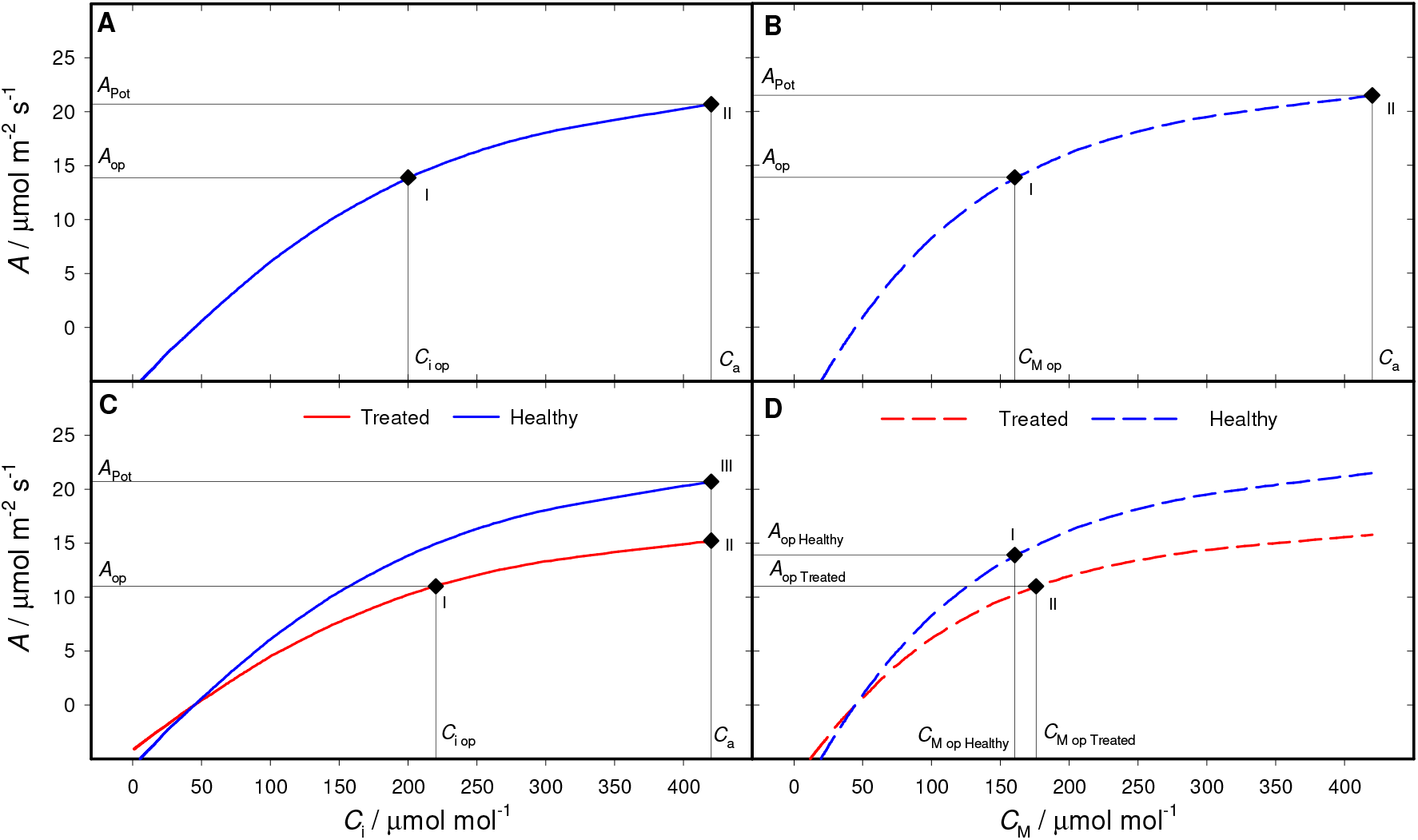
Quantifying limitations and contributions to changes in photosynthesis Stomatal limitation. Panel **A** shows a photosynthetic model describing the dependence of assimilation (*A*) upon CO_2_ concentration in the substomatal cavity (*C*_i_, solid line, Eqn 9) of healthy plants (blue). In this example with hypothetical data, operational conditions correspond to an ambient CO_2_ concentration (*C*_a_) of 420 μmol mol^-1^ and a fixed light intensity (the level is irrelevant here). It attains a rate of assimilation (*A*_op_) of 13.9 μmol m^-2^ s^-1^, and has a CO_2_ concentration in the substomatal cavity (*C*_i op_) of 200 μmol mol^-1^, corresponding to an operational stomatal conductance to CO_2_ 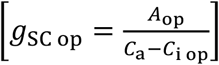of 0.063 mol m^-2^ s^-1^ (Table 2). In the transition between operational State I and hypothetical State II assimilation is driven to its potential as if intercellular spaces were directly exposed to external CO_2_ concentration, *C*_a_ (Eqn 9 calculated for *C*_i_ = *C*_a_). Assimilation increased from 13.9 to 20.7 μmol m^-2^ s^-1^; the total contribution of *C*_i_ to the change in assimilation [*Ξ*(*C*_i_)_*A*_] was 6.9 μmol m^-2^ s^-1^; the relative contribution of *C*_i_ to the change in assimilation [ρ(C_i_)_*A*_] was 1 because *C*_i_ was the only input to vary between the two states; the limitation imposed by *C*_i_ to assimilation [*L*(*C*_i_)_*A*_] corresponds to stomatal limitation, was 0.33. *S*(Ω)_*a*_, and *η*(Ω)_*a*_ are shown in Sheet 1 of Workbook III, calculated for each step of the transition.

##### Diffusional limitation

The analysis of diffusional limitation quantifies the total effect of diffusion from atmosphere to the sites of carboxylation, and requires *g*_M_. This not straightforward to estimate and involves combined measurements of gas exchange and variable fluorescence in C_3_ plants (Bellasio et al. 2016b), or isotopic methods in C_3_ and C_4_ plants (Ubierna et al. 2017; Busch et al. 2020). Diffusional limitation is evaluated in the same way as stomatal limitation, using *C*_M_ in place of *C*_i_. Practically, for each measured value of *C*_i_, the corresponding value of *C*_M_ is calculated as 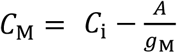 . Eqn 9 is fitted as described above, this time to the *A*/*C*_M_ response curve obtaining new *C*_M_-based *CE*, and *ω* (Table 2). Eqn 11 is calculated in all *K* intervals between state I where *g*_S_ and *g*_M_ take the operational values and, and state II, the hypothetical scenario where *g*_S_ and *g*_M_ are high. Application is exemplified in Figure 3b and the corresponding calculations are in Sheet 2 of Workbook III.

##### Stomatal and non-stomatal limitation

Evaluating the stomatal limitation of treated plants using the procedure shown above (transition number 5 in Figure 1b) requires knowing the *A*/*C*_i_ responses of the treated plant. When these are impractical or impossible to measure, their parameters need to be estimated. In the method that I recently proposed (Bellasio 2023), the *A*/*C*_i_ curve of the treated plant is obtained by ‘flattening’ the *A*/*C*_i_ curve of the healthy plant (red line in Figure 2C) to match the pair (*C*_i op_, *A*_op_) measured for the treated plant. The assumptions made are that the curvature *ω* and the CO_2_ compensation point Γ are not affected by the treatment, while the horizontal asymptote *A*_SAT_ and the initial slope *CE* maintain their ratio *μ* = *A*_SAT_/*CE* constant. Eqn 9 is solved for *CE* as:

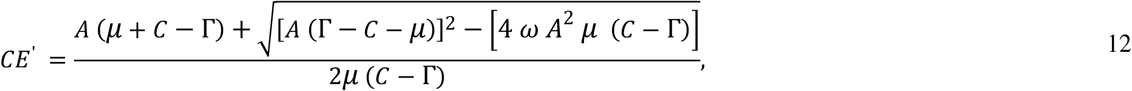

*CE*′, the estimated value of *CE* of the treated plant, is directly calculated by inputting measured *A*_op_ and *C*_iop_ or *C*_Mop_ in Eqn 12; *A*_SAT_′, the estimated value of *A*_SAT_ in the treated plant, is, by assumption, *μ CE*′. The difference between the potential attained by the treated plant and that of the healthy plant is conventionally referred to as non-stomatal limitation, which can be evaluated in a second transition (number 6 in Figure 1b). The procedure is shown in Figure 3C and the corresponding calculations and control analyses are shown for each step in Sheet 3 of workbook III.

An alternative procedure, proposed here anew, lifts limitations in a single transition between the operational conditions of the treated plant and the potential conditions of the healthy plant (number 7 in Figure 1b).

If *A*/*C*_i_ curves and *g*_M_ are available for both the healthy and the treated plants, diffusional limitation can be resolved in its mesophyll and stomatal components. A first transition may lift diffusional limitation (number 8 in Figure 1b), while a second transition may lift non-diffusional limitation, as for non-stomatal limitation (number 6 in Figure 1b). A single transition lifting diffusional and non-diffusional limitation concurrently is also possible. Also these two alternatives are new to this study.

##### Stomatal, mesophyll and non-diffusional contributions

If *A*/*C*_i_ curves and *g*_M_ are available for both the healthy and the treated plants, the effect of the treatment can be resolved in its biochemical, mesophyll and stomatal components in a single transition (number 4 in Figure 1b). This is essentially a simple generalisation of the method of Buckley and Diaz-Espejo (2015). A numerical example is shown in Figure 3D, and the corresponding calculations and control analysis are in Sheet 4 of Workbook III. If *g*_M_ for the treated plant is not available, the corresponding transition (number 3 in Figure 1b) can be implemented in with the same procedure, using the *C*_i_-based parameterisation, and by setting an arbitrarily high value for *g*_M_. This variant is new to this study.

#### Control analysis

The extent to which assimilation is controlled by a marginal change in the underpinning variables (typically, *g*_S_ and *g*_M_ alongside other inputs in ***b***) is quantified by sensitivity. The strength of this analysis lies in being univocal in any given state, allowing for independent evaluation of plants (*e*.*g*. healthy plants under ordinary operational conditions) and *a posteriori* comparison (*e*.*g*. across species in the phylogeny (Gago et al. 2019)).

This analysis can be implemented using the logic of previous examples by imagining that the transition between states becomes arbitrarily small. In practice, Eqn 11 is used calculated for the state of interest, and that in which *g*_S_ and *g*_M_ (alongside *A*_SAT_, *CE*, *ω*, and Γ, or a subset thereof) differ by a marginal increment (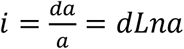, for *i*→0). Calculations for the healthy plant (Table 2) with *i*=0.0001 applied to *g*_S_, *g*_M_, *A*_SAT_, and *CE* are included in Sheet 7 of Workbook III. The key outputs are absolute and relative sensitivity (for instance, 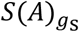 averaged 113 μmol mol^-1^, 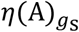 averaged 0.5), and the relative change in assimilation, which adds to unity when compounding all inputs (in the example, 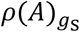was 0.5 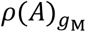 was 0.09, 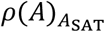 was 0.17, and p(A)_CE_ was 0.25). In small transitions, 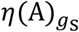 and 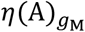 correspond to *l*_S_ and *l*_M_ calculated after Grassi and Magnani (2005) while *L*(*g*_S_) and *L*(*g*_M_) correspond to *S*_L_ and *MC*_L_ in their Eqn 6.

#### Light and non-light restrictions

##### Light limitation

Light limitation analysis can be utilized to quantify the reduction of assimilation due to shading, relative to a hypothetical case where the leaf would be exposed to maximum lighting. Similar to the procedure used for stomatal limitation analysis, light response curves are measured under specific growth conditions (CO_2_ concentration, temperature, *etc*.). Then, the dependence of assimilation on light intensity is described with the model of Prioul and Chartier (1977) after Bellasio et al. (2016b) as:

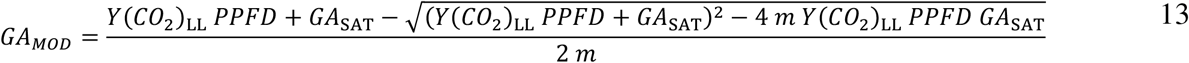

Eqn 13 (coded in the ‘GAvPPFD’ Sheet of Workbook I) is fitted to the experimental data to derive the parameters for the healthy plant. Light limitation is quantified by evaluating a single transition between operational conditions, and where light limitation is assumed to be absent (number 1 in Figure 1c), using the procedures described in Workbook II. A worked example is provided in Sheet 6 of Workbook III.

##### Light contribution

The previous procedure is apt for analysing light contributions between two real conditions of interest by specifying the second condition in the parameterization of State II (number 2 in Figure 1c).

##### Light and non-light limitations

The analysis of light and non-light limitations can be used to distinguish the effect on assimilation directly due to shading from that due to the downregulation of the biochemical potential that occurred in a treated plant. Analogous to the approach used to resolve diffusional and non-diffusional limitations, two variants are possible. In the first case, light limitation is lifted first (number 3 in Figure 1c), and the remainder is the difference with the potential of the healthy plant (number 4 in Figure 1c). Alternatively, limitations are lifted in a single transition (number 5 in Figure 1c, exemplified in Sheet 7 of Workbook III).

### Mechanistic models

The empirical models shown above are suitable for most applications, including when parameters of mechanistic models are available from previous experiments. Empirical parameters can be estimated approximately: *CE* corresponds to 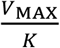(where *V*_MAX_ may be Rubisco or PEP carboxylase maximum rates and *K* is the corresponding Michaelis-Menten constant, adjusted for oxygen concentration in Rubisco’s case); 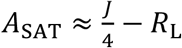, where *J* is the rate of electron transport, and *R*_L_ is light respiration, or, for C_4_ plants 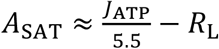, where *J*_ATP_ is the rate of ATP production; Γ is typically available in the literature (alternatively, it can be calculated from *R*_L_ and *K* using, for instance, Eqn 2.38 or Eqn 4.50 in von Caemmerer (2000) for C_3_ plants or C_4_ plants, respectively); whilst the curvature *ω* can typically be assumed as 0.7. For a more precise analysis, it is recommended to generate synthetic *A*/*C* response curves using 10-15 values of *C*_i_ (or *C*_M_) as inputs for the same mechanistic model formulations from which the mechanistic parameters were originally derived. These synthetic *A*/*C* curves can then be used to derive empirical parameters through curve fitting, and then used to evaluate restrictions, as described above.

Specific applications may require direct use of mechanistic models in restriction analyses. Mechanistic models are typically more complicated than empirical, rely on a number of assumptions, are valid only when assumptions hold (*e*.*g*. a C_4_ model is not meaningful for mechanistically describing C_3_ plants) and they are less flexible in data fitting. In general, they are recommended only for cases where a description of the processes is specifically required. For instance, they can resolve non-diffusional restrictions, such as distinguishing between those due to light reactions or carbon metabolism. Well-thought-out procedures are needed, designed specifically to address the experimental question.

The method presented by Buckley and Diaz-Espejo (2015) differs from the analysis of mesophyll, stomatal, and non-diffusional contributions exemplified above only in that it uses an update of the C_3_ model by Farquhar et al. (1980). This model has previously been used for the analysis of limitations imposed by Rubisco activation in its enzyme limited submodel (Deans et al. 2019) or in the full formulation (Taylor and Long 2017), and can also be used to quantify limitations imposed by electron transport and triose phosphate utilisation (Buckley and Diaz-Espejo 2015). However, this method is less widely applicable than the empirical procedure described above: it is meaningful only for C_3_ plants; obtaining specific values for the Rubisco Michaelis-Menten constants for carboxylation and oxygenation (*K*_C_ and *K*_O_) for measured leaves may not be readily achievable, and the cut-off point for enzyme and light limitation must be assumed.

A C_4_ mechanistic model may serve for the quantification of restrictions due to diffusion through stomata or the mesophyll, as well as restrictions caused by other factors like light intensity, the light-saturated rate of ATP generation (*J*_SAT_), the curvature of the light dependence of ATP generation (θ), the initial quantum yield for ATP generation *Y*(*J*_ATP_)_LL_, respiration in the light, Rubisco and phosphoenolpyruvate carboxylase CO_2_ saturated rate of carboxylation (*V*_CMAX_ and *V*_PMAX_), the factor that partitions ATP between PEP regeneration and the remainder of photosynthetic activity (*x*), and bundle sheath conductance (*g*_BS_). For example, it can be used to determine the contribution of light versus that of *g*_BS_ in the reduction of assimilation for plants grown under shading (Bellasio and Griffiths 2014b). For a corrected formulation refer to Bellasio et al. (2017).

For simulating more complex scenarios and addressing a broad range of questions relating carbon and light reactions I recommend utilizing the Bellasio and Farquhar (2019) and Bellasio and Ermakova (2022) analytical models. The analytical model of Bellasio and Farquhar (2019) allows the quantification of limitations imposed by a wide range of real or imagined scenarios, in any photosynthetic type (C_3_, C_2_, C_2_+C_4_, C_4_), intermediates thereof, and engineered transitions between them, stomatal opening and closure, different stomatal responsiveness to biochemical forcing or drought, and a wide range of characteristics of the electron transport chain (*e*.*g*. shifting the ratio between ATP and NADPH production by modifying the fraction of cyclic electron flow or the NDH pathway, *etc*.). The analytical model of C_4_ photosynthesis of Bellasio and Ermakova (2022) relates to leaf anatomy following Bellasio and Lundgren (2016) and explicitly accounts for light harvesting partitioning between cell compartments. It was specifically developed for studying differences between C_4_ photosynthetic subtypes, and maintains validity at very low irradiance. It features two electron transport chains, C_4_ cycle reactions, and C_3_ cycle reactions accommodating a variety of possibilities for parameterisation.

## Discussion

I set out to generalise the analyses of photosynthetic restrictions, providing a set of definitions that integrate all common and several new procedures, which I illustrated using synthetic data.

The new framework is general, as it is valid for all photosynthetic types and models, and it comprehensively quantifies limitations, contributions, and photosynthetic control (sensitivity and elasticity) within a single procedure. This represents a significant advancement since the most widely used protocols – control analysis (Grassi and Magnani 2005), contribution analysis (Buckley and Diaz-Espejo 2015) and limitation analysis (Farquhar and Sharkey 1982) – were traditionally independent. This method is formulated using the principles of Buckley and Diaz-Espejo (2015), and I demonstrated its convergence with Farquhar and Sharkey (1982) and Grassi and Magnani (2005) for small transitions (see ‘Control analysis’ section). For complete generalisation, I also formulated within this framework the method of Björkman et al. (1980), which, although widely used, has never been formalised (Figure S1a, and discussed below).

When only one quantity varies, procedures are unambiguous. This is the case for diffusional limitation and light limitation analyses, which are evaluated by formulating calculations so that only one quantity (variable *a* in Eqn 1) varies between transitions, while the other inputs (vector ***b*** in Eqn 1) remain constant. In all those cases the method is equivalent both for limitation and contribution analysis. When multiple quantities vary, methodological variants are possible (the number of variables changing raised to the power of the number of transitions), but choices are straightforward or alternatives are interchangeable, as I will now discuss.

In contribution analysis, all quantities can be presumed to change concurrently and linearly between their initial and final values (number 3 in Figure 1b), as proposed by Buckley and Diaz-Espejo (2015), thus there is no longer ambiguity in the path to follow. In classical literature, it had been proposed to assume that stomata respond before any change in leaf biochemical processes occurs, or the reverse. The discrepancy between paths was often large, for instance Assmann (1988) reported a five-fold difference. I calculated both path dependent alternative (shown in Figure S1b and calculated in sheets 9 and 10 of Workbook III), and found that the discrepancy for stomatal contribution was relatively small about ±20% of the independent path. Yet, path dependent methods are not recommended. While their assumptions may partially hold under specific conditions, such as when stomatal adjustment is much slower than photosynthetic activation (Deans et al. 2019), they are generally invalid. In fact, during the imposition of a treatment, photosynthetic characteristics typically respond progressively, driven by a common factor – the treatment – and are therefore not independent.

In the analysis of limitations, there is the possibility to opt for a single or two-tiered transition. The distinction is merely conceptual, as the results do not differ. Much of the past debate has been driven by methodological papers prioritizing the breadth of alternative approaches rather than narrowing their choice. In contrast, focusing on the experiment and its research questions will naturally determine the appropriate method to follow. Plants are often subject to permanent or mid-term acclimation treatments (*e*.*g*., differing nitrogen levels, watering levels, growth light conditions, growth CO_2_ levels, soil types, fertilisation, mycorrhization, *etc*.), or differ by any genetic trait, whether natural or engineered. These traits are not subject to short-term changes, and it is possible to devise a hypothetical scenario whereby limitations are lifted independently, assuming the characteristics of the plant remain constant. Applying this principle involves evaluating two transitions (numbers 5 and 6 in Figure 1b), which is the conventional approach used in the literature. I presented a novel single-transition variant for limitation analysis (number 7 in Figure 1b), which has the benefit of being univocally defined. Under current assumptions − that inputs scale linearly (Eqn 2) and diffusional limitations are fully lifted in the idealized state, where the total conductance to CO_2_ becomes high – the single-transition variant accurately tracks the two-transition variant, without differences in results. (This can be verified by typing 1 in cell C1 and “n” in cell F10 in Sheet 3 of Workbook III) Therefore, unless additional assumptions are introduced, they can be used interchangeably.

Sometimes it may be difficult to determine the characteristics ***b*** of the treated plants. Through an elegant experimental routine, Taylor and Long (2017) measured rapid *A*/*C*_i_ responses in real time during the treatment’s course, using modern gas exchange equipment and routines (Stinziano et al. 2017). However, often this is impractical; in which case the characteristics of treated plants must be estimated. A simple and widely used method is that of Björkman et al. (1980), which the estimation follows the assumption that lifting stomatal limitation in the treated plant would result in the same increase in assimilation as in the healthy plant (details and mathematical formalisation in Figure S1a). This simplification is generally acceptable for C_4_ plants because their *A/C*_i_ curves are flat for *C*_i_ > *C*_iop_. For C_3_ plants, it may be acceptable only when the treatment effect is small, but stomatal limitation is overestimated when the assimilatory potential is severely reduced by the treatment. The method I followed here to derive an analytical solution is based on Bellasio (2023). It involves flattening the *A/C*_i_ curves of treated plants, under the assumption that *CE* and *A*_SAT_ maintain their ratio (Eqn 12). This corresponds mechanistically to fixing the *J*/*V*_MAX_ ratio, and may not have general validity, as the treatment’s effect on parameters may be species- and treatment-dependent. The coded procedure allows to input the actual ratio, which should be preferred if available. Γ and ω are maintained invariant based on the rationale that they are underpinned by enzymatic properties (see ‘Mechanistic models’ above), and indeed measurements show that, for instance, they did not significantly change in both C_3_ and C_4_ plants during seasonal drought (Bellasio et al. 2022).

The framework avoids two erroneous practices: that of approximating curves with lines in finite intervals (see Introduction), and that of overestimating their slopes. The latter practice was introduced by Tomás et al. (2013) and gained traction in later studies, for instance by Tosens et al. (2016); Carriquí et al. (2015); Carriquí et al. (2019); Perera-Castro et al. (2022); Hu et al. (2024). The slope of the *A*/*C*_M_ response curves required for the linear approximation of the infinitesimal analysis of Grassi and Magnani (2005) was derived under low CO_2_ concentrations – where it is typically several-fold higher than under ordinary operational conditions. This inevitably leads to largely overestimating the importance of stomata and mesophyll diffusion – and has likely brought to overstating their importance – in restricting assimilation.

The analysis of light and non-light restrictions is original to this study. Lifting light limitation may not involve exposing leaves to infinite light intensity but just to full sunlight, making it difficult to pinpoint the real versus hypothetical conditions, creating confusion between contribution and limitation analyses. A decrease in assimilation may result from a combination of direct and indirect effects of shading, such has the decrease in biochemical potential that occurs in a leaf shaded by the overgrowth of a new canopy (Bellasio and Griffiths 2014a). Because these responses are not independent, in this case the analysis of contributions in a single transition may be appropriate. Alternatively, if the treatment has affected the biochemical potential independently of light, such as in cases of pathogen attack or nitrogen level changes (Dewar et al. 2012), then limitation analysis with a two-tiered transition may be preferable.

## Conclusion

I urge the community to critically reconsider the relative importance of stomatal, mesophyll, and non-diffusional restrictions, especially when they were evaluated using the linear approximation in finite intervals by Grassi and Magnani (2005), and the initial slopes of the *A*/*C*_M_ curves by Tomás et al. (2013), which are erroneous.

## Supporting information

Example of calculations

Curve fitting

Restriction analysis

## Availability

The workbooks coding curve fitting, procedures for restriction analysis and worked examples are available in the Supporting Information of this article. A curated version is available on GitHub at the link [https://github.com/chandrabellasio/Quantifying-Photosynthetic-Restrictions-]. The C_3_ and C_4_ models of Farquhar et al. (1980) and Berry and Farquhar (1978) in the formulations of Bellasio et al. (2017) can be made available upon request. Spreadsheets coding the Bellasio and Farquhar (2019) and Bellasio and Ermakova (2022) analytical models are available in the Supporting Information of the original publications.

## Acknowledgments

This work slowly emerged from the requests I received during the four years of the gestation of my previous commentary (Bellasio 2023), eventually expanding into a standalone paper. Reaching this point would have been impossible without the trust of Editor Patrick Morgan and the patience of the reviewers, who managed to navigate my early drafts and provide the suggestions that shaped this research at various stages. I am grateful to Alessio Fini, Hilary Stuart-Williams, Ross Deans, and Jaume Flexas for their valuable discussions.

I have no competing interests.

## Funding

I gratefully acknowledge funding by a Science Foundation Ireland Individual Pathway Fellowship (21/PATH-S/9322), and by a European Union’s Horizon 2020 research and innovation programme MSCA individual fellowship (grant agreement ID No: 702755).

Plants were either untreated or subjected to a hypothetical treatment which reduced the photosynthetic potential. The rates of assimilation (*A*) and CO_2_ concentration in the substomatal cavity (*C*_i_) attained in the conditions in which plants were grown are denoted as operational. Gas exchange measurements at the lab bench determined the responses of *A* to *C*_i_. Values of CO_2_ concentration at the sites of carboxylation was calculated using mesophyll conductance values of

0.35 mol m^-2^ s^-1^ and 0.25 mol m^-2^ s^-1^, previously obtained for healthy plants and treated plants, respectively. Assimilation is expressed in μmol m^-2^ s^-1^, CO_2_ concentration in μmol mol^-1^. Non-rectangular hyperbolas were fitted either to the *A*/*C*_i_ or to *A*/*C*_M_ curves to obtain two sets of fitted parameters reported at the bottom using the fitting tool of Bellasio et al. (2016b), copied in the sheet ‘AvCi’ of the workbook provided in Supporting Information for convenience.

### Diffusional limitation

Panel **B** shows the modelled dependence of assimilation (*A*) upon CO_2_ concentration in the mesophyll (*C*_M_, dashed line, Eqn 9) of healthy plants. Diffusional limitation is quantified by evaluating transitions from the operational conditions of state I with physiological *g*_M_ and *g*_S_, to a state II where the hindrance to CO_2_ diffusion caused both by the stomata and by the mesophyll is removed by increasing *g*_M_ and *g*_S_ to high values. Assimilation increased from 13.9 in state I, to 21.5 μmol m^-2^ s^-1^ in state II; diffusional limitation was 0.36; details are in Sheet 2 of workbook III.

### Stomatal and non-stomatal limitation

Panel **C** shows the modelled *A*/*C*_i_ curves of healthy (blue) and treated (red) plants. State I is the operational conditions of the treated plant (diamond ‘I’). State II is where, with otherwise invariant parameterisation, the stomatal barrier is removed by increasing *g*_M_ and *g*_S_ so that *C*_i_ reaches approximately the value of *C*_a_ of 420 μmol mol^-1^. State III is where *C*_a_ is invariant, while the parameters *A*_SAT_ and *CE* are driven to those of the reference plant. The treated plant had an *A*_op_ of 11 μmol m^-2^ s^-1^, and a *C*_i op_ of 220 μmol mol^-1^, the healthy plant is the same (Table 2). In the transitions, assimilation increased from *A*_op_ of 11 in state I, to *A*_Ca_ of 15.2 μmol m^-2^ s^-1^in state II to *A*_Pot_ of 20.7 μmol m^-2^ s^-1^ in state III. Stomatal limitation, *L*(g_S_)_*A*_, was 0.20; the sum of the other limitations (*A*_SAT_, *CE, ω*, and Γ), that is, non-stomatal limitation, was 0.26. Calculations are shown in Sheet 3 of Workbook III.

### Contribution analysis

Panel **D** shows the modelled *A*/*C*_M_ curves of healthy (blue) and treated plants (red). Healthy plants’ *A*/*C*_M_ responses, *A*_op_, *C*_M op_, *g*_SC op_ and *g*_M_ are from previous examples while *g*_SC op_′ was 0.055 mol m^-2^ s^-1^. Parameters of the fitted to *A*/*C*_M_ responses are in Table 2. State I and II represent generic operational points of healthy and treated plants, respectively. Stomatal, mesophyll and non-diffusional contribution are quantified in a single transition whereby quantities change linearly between values of healthy and treated plants. The total diffusional contribution was -0.9 μmol m^-2^ s^-1^, resolved in a stomatal contribution of -0.5 μmol m^-2^ s^-1^, and a mesophyll contribution of -0.4 μmol m^-2^ s^-1^. The total non-diffusional contribution was -1.7 μmol m^-2^ s^-1^. The calculations are in Sheet 4 of Workbook III.

---

**Figure S1.**
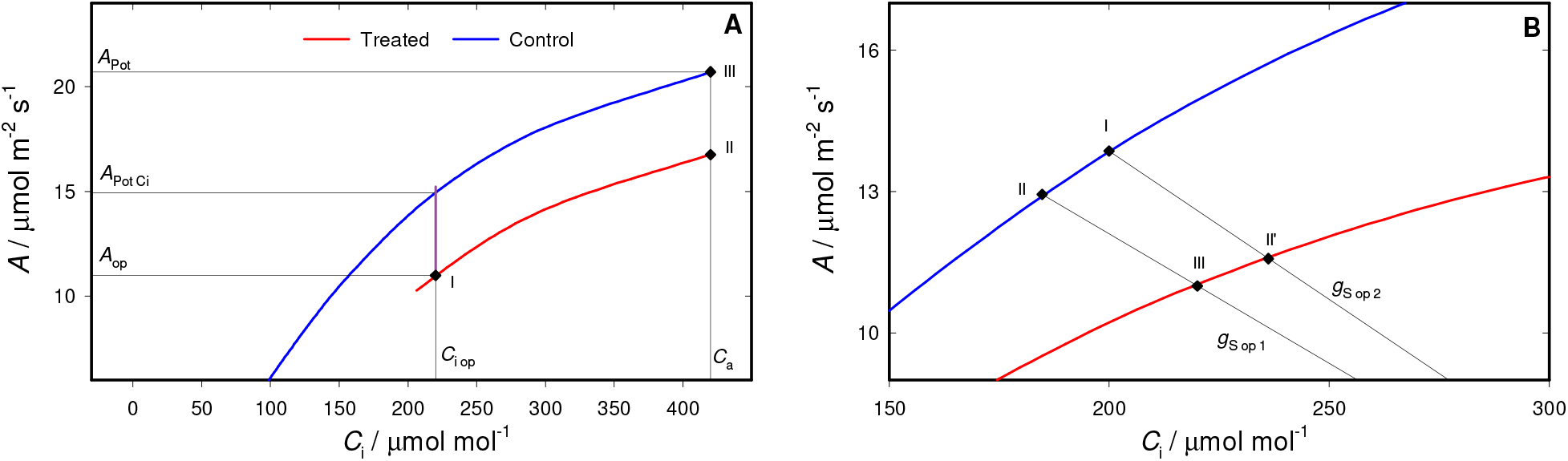
Alternative path dependent methods. Panel **A** shows the method of Björkman et al. (1980). The *A*/*C*_i_ curve for the treated plant (in red) is approximated with that of the healthy plant (in blue) for *C*_i_ > *C*_i op_, translated vertically to match the operational assimilation of the measured plant as:

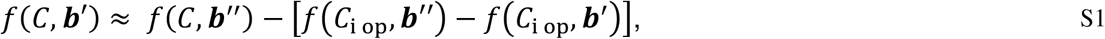

where ***b***′ and ***b***′′ are the set of parameters of the treated, and healthy plant, respectively; *f*(*C*, ***b***′) is the *A*/*C* function to be estimated; *f*(C, b′′) is the known *A*/*C* function with the parameters ***b***′′ determined for the healthy plant; the expression in square brackets is the down-translation (Figure 4A). *A*_op_ is the rate of assimilation of the treated plant in operational conditions, *f*(*C*_iop_, b^i^) in Eqn S1; A_Pot Ci_, is the rate of assimilation that the healthy plant would have if *C*_i_ were that of the treated plant in the operational condition *C*_iop,_, *f*(*C*_iop_, ***b***′′) in Eqn S1; *A*_Pot_ is that a healthy plant would have in absence of stomatal barrier, *f*(*C*_a_, ***b***′′) in Eqn S1. Limitations are evaluated by removing stomatal limitation first in the transition between state I and state II, and then by removing non-stomatal limitation in the transition between state II and state III. The conventional three point calculation (Bellasio et al. 2023) gives the same results as the current framework: the total stomatal contribution Ξ(*C*_i_)_A_ is *A*_Pot_ − *A*_Pot Ci_, and the total non-stomatal contribution Ξ(*L*_NS_)_*A*_ is *A*_Pot Ci_ − *A*_op_. Calculation of Eqn S1 for each step of the first transition is shown in Sheet 8 of Workbook III. Panel **B** shows two path-dependent variants of contribution analysis where a healthy plant (in blue) undergoes a generic treatment that restrict assimilation (in red). State I and III represents a generic operational point of the treated plant, and healthy plant, respectively, while state II is an intermediate state through which the transition is assumed to occur. The first path assumes that in response to the treatment stomata respond before any change in leaf biochemical processes occurs, that is, a transition from state III to state II, with reduced *g*_S_, and then to state I, with lower photosynthetic potential but equal *g*_S_. The second variant assumes that biochemical regulation occurs before stomata respond, that is, a path through state II′ where *g*_S_ equals that of the healthy plant, while the parameterization corresponds to that of the treated plant. In the path ‘stomata first’ during the transition between state I and state II, Ξ(*g*_*S*_)_*A*_ was -1.1 μmol m^-2^ s^-1^. In the path ‘mesophyll first’ during the transition between state II and state III, Ξ(*g*_*S*_)_*A*_ was -0.7 μmol m^-2^ s^-1^. The calculations are in Sheets 9 and 10 of Workbook III.

